# Medial prefrontal cortex input to lateral entorhinal cortex supports both encoding and retrieval of associative recognition memory

**DOI:** 10.1101/2025.08.14.670409

**Authors:** Lisa Kinnavane, Gareth RI Barker, Paul J Banks, Zafar I Bashir, Elizabeth Clea Warburton

## Abstract

Associative recognition memory allows us to form representations of items and their environment and to judge the novelty of such representations. This memory is dependent on a brain circuit that includes interactions between medial prefrontal cortex (mPFC) and lateral entorhinal cortex (LEC); however it is unknown whether the interaction of these brain areas is required for memory encoding, retrieval or both processes. Furthermore, little is known as to whether indirect or direct mPFC-LEC connections are critical for associative recognition memory and, if the latter, in which direction information travels. To address these questions, we first performed pharmacological disconnection of mPFC and LEC, finding that mPFC-LEC interaction is required for both memory encoding and retrieval. Next, we optogenetically inhibited projections from mPFC to LEC, showing that this projection was crucial for both encoding and retrieval of both object-in-place and object-in-context recognition memory when a 1 h, but not a 5 min, memory retention delay was used. These data show that a direct connection from mPFC to LEC is critical for associative recognition memory, in a delay-dependent manner.

## Introduction

Associative recognition memory enables the recognition of individual items in relation to the location of other items with which they co-occur, or the context in which they were last encountered. Such memory contributes to episodic memory formation (Endemann and Kamp, 2025) and is crucial for navigation and foraging. Concerning the neural structures which underpin associative recognition, studies have revealed that such memory is significantly impaired in patients with medial temporal lobe (MTL) or frontal cortex damage (Janowsky et al., 1989; Duarte et al., 2005; Bastin et al., 2014; Hampstead et al., 2018) including in patients with dementia (Lee et al., 2003; Blackwell et al., 2004). Further, lesion studies in animals suggest that the hippocampus (HPC) and medial prefrontal cortices (mPFC) are critical for associative object recognition memory (Bussey et al., 2000; Barker and Warburton, 2011) and further that functional interactions between these regions underpin both encoding and retrieval (Barker and Warburton, 2015). Thus fronto-temporal networks may play a key role in associative recognition memory.

The lateral entorhinal cortex (LEC) is reciprocally connected with the PFC (Conde et al., 1995; Delatour and Witter, 2002; Vertes, 2004; Hoover and Vertes, 2007). In rodents, selective lesions in the LEC have been shown to disrupt associative recognition memory tasks such as object-in-place (OiP) or object-in context (OiC), in which recognition of a familiar object in a novel environment or context is tested (Hunsaker et al., 2013; Wilson et al., 2013a; Kuruvilla et al., 2020). In addition, performance of an OiC task has been shown to increase expression of the immediate early gene c-fos in LEC (Wilson et al., 2013b). Thus it has been hypothesised that fronto-entorhinal networks are critical for associative recognition memory, in support of this disconnection lesions of mPFC and LEC impaired associative recognition memory formation (Chao et al., 2016). However it is not known if this fronto-entorhinal network is critical for both memory encoding and retrieval and whether direct or indirect connections play a critical role in its function.

Recently we have begun to map how specific pathways within the associative recognition memory network differentially mediate the separate phases of memory encoding and memory retrieval. For example, we have shown that mPFC afferents from the nucleus reuniens (Barker et al., 2021) are selectively required for encoding but not retrieval, whilst efferents to nucleus reuniens (Barker et al., 2021) or medial dorsal thalamus (Culleton et al., unpublished) are selectively required for retrieval but not encoding. Until now such observations have been restricted to mPFC-thalamic connections, and the contribution of mPFC outputs to cortical regions, such as the LEC have not been investigated. Furthermore, the LEC and mPFC are highly interconnected both directly and indirectly (e.g. via the HPC, or nucleus reuniens of the thalamus) (Agster and Burwell, 2009; Dolleman-van der Weel et al., 2019). Thus, here we examined the role of connections between mPFC and LEC in encoding compared to retrieval for both an OiP and OiC task firstly by disconnection analysis. We then determined the contribution of the direct mPFC to LEC projection at both short (5min) and longer delays (1h) using an optogenetic inhibition approach.

## Materials and Methods

### Subjects

All experiments were carried out in naïve male Lister Hooded rats (Envigo) weighing 350 – 450 g at the start of the experiments. Experiment 1 used 12 rats and experiment 2 used 24 rats. Animals were housed in groups of 2-4, under a 12 h light/dark cycle (light phase, 8.15 P.M. to 8.15 A.M) with ad libitum access to food and water. Behavioural testing was conducted during the dark phase of this cycle. All animal procedures were performed in accordance with United Kingdom Animals Scientific Procedures Act (1986) and associated guidelines. All efforts were made to minimise any suffering and the number of animals used.

### Surgical procedures

#### Cannulae implantation in medial prefrontal cortex (mPFC) and lateral entorhinal cortex (LEC) (Experiment 1)

Rats were anesthetized (isoflurane: induction 4%; maintenance 2%). The scalp of the animals was shaved before they were placed in a stereotaxic frame (Kopf Instruments, USA) with the incisor bar set to achieve a flat skull (approximately 3.3-mm below the interaural line). Eye drops (0.1% sodium hyaluronate; Hycosan, UK) were applied. The scalp was further anesthetized using lidocaine (5% m/m; TEVA; UK) and disinfected with chlorhexidine, cut, and retracted. Rats were implanted with a double cannula targeted to bilateral mPFC and two single cannulae targeted to bilateral LEC (Figure 1A). For the mPFC; the stainless-steel double guide cannula (26 gauge; centre to centre distance 1.5mm; Plastics One) was implanted through burr holes in the skull at the following coordinates relative to skull at bregma: anterior–posterior (AP) +3.2 mm, mediolateral (ML) ±0.75 mm, lowered 3.5 mm relative to surface of the skull. For the LEC sites, a single stainless-steel guide cannula (26 gauge; Plastics One) was implanted bilaterally through a burr hole in the skull at the following coordinates relative to skull at bregma: AP –6.4 mm, ML ±4.5 mm and lowered 7.7 mm relative to surface of the skull with the manipulator arm at an angle of 12° to the vertical. The cannulae were anchored to the skull by stainless steel skull screws (Plastics One) and dental acrylic. After surgery, each animal was given fluid replacement therapy of at least 5 ml of glucose saline subcutaneously and analgesia intramuscularly (0.05 ml of 0.3 mg/ml buprenorphine). Animals were housed individually for 7 days to recover from surgery and in pairs thereafter. The animals recovered for at least 14 days before habituation to the testing arena began. Between infusions, 33-gauge obdurators (Plastics One) were used to keep the cannulae patent.

**Figure 1.**
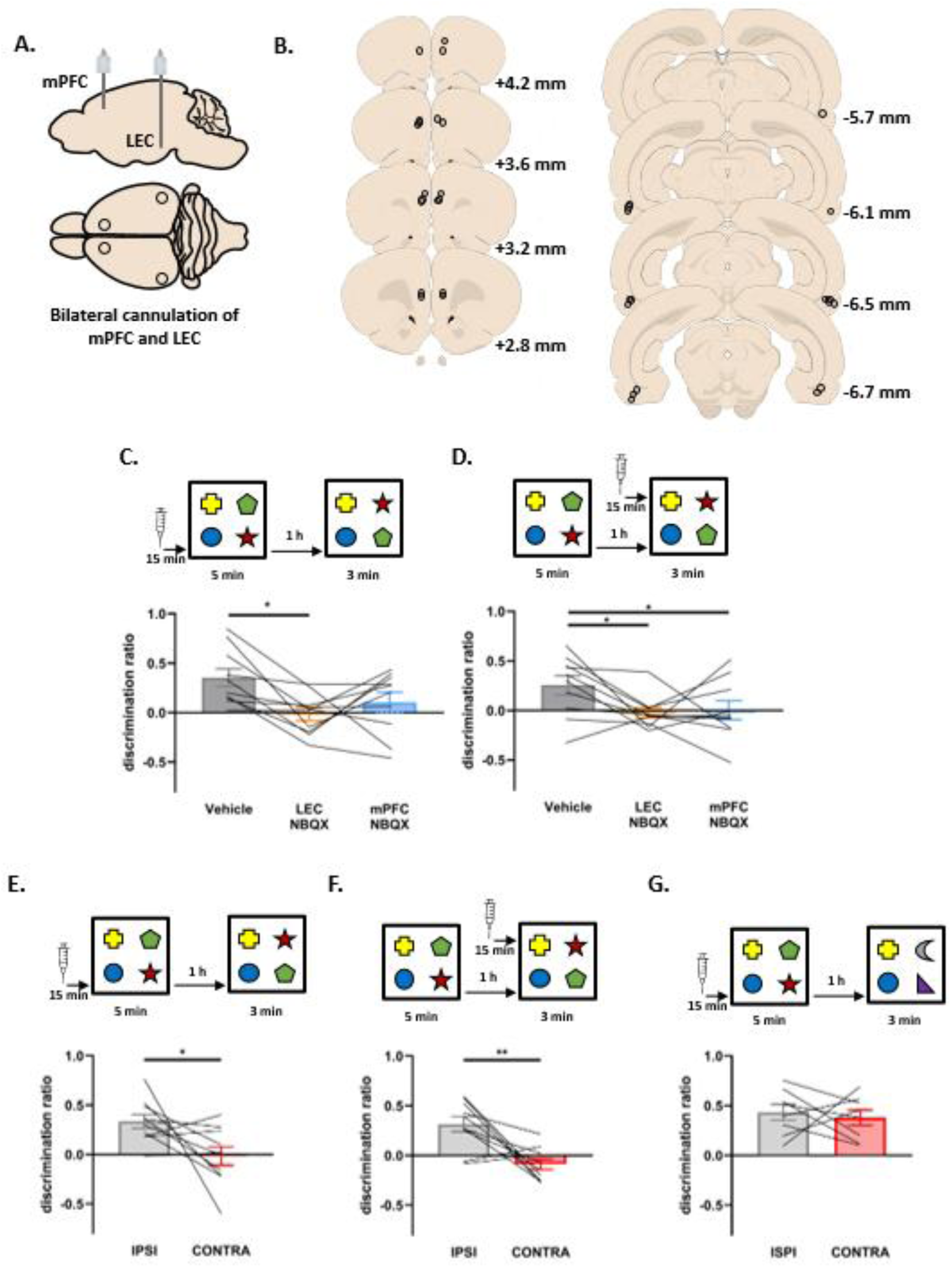
mPFC-LEC disconnection impairs both encoding and retrieval of object-in-place memory. **A**: Schematic of bilateral cannulation of mPFC and LEC. **B:** Post-hoc reconstruction of cannula placement. Numbers denote distance from bregma in mm. **C:** Effect of bilateral infusions prior to sample phase. **D:** Effect of bilateral infusions prior to test phase. **E:** Contralateral, but not ipsilateral NBQX infusion prior to sample phase impairs OiP memory. **F:** Contralateral, but not ipsilateral NBQX infusions prior to test phase impairs OiP memory. **G:** Novel object preference is not impaired by mPFC-LEC disconnection.

#### Inhibitory opsin viral injections in mPFC and optic fibre implantation in LEC (Experiment 2)

Optogenetic transduction of mPFC neurons was achieved as described previously (Kinnavane and Banks, 2022) using AAV5-CaMKii-eARCH3.0-eYFP (UNC Vector Core; 3.4 × 10^12^ genome copies/mL) and in the control group AAV5-CaMKii-eYFP (UNC Vector Core; 3.6 × 10^12^ genome copies/mL). Rats were anesthetized as described above and secured in a stereotaxic frame with the incisor bar set 3.3-mm below the interaural line. Injections were targeted to the mPFC; bilateral burr holes were made in the skull at the following coordinates with respect to bregma: AP +3.1 mm, ML ± 0.7 mm. Virus was loaded into a 33-gauge 12° bevelled needle (Esslab) attached to a 5 µL Hamilton syringe with the eyelet of the needle facing posteriorly. The needle was lowered 4.5-mm below the surface of the skull measured from the burr hole and 2 µL of virus was delivered via each burr hole at a rate of 200 nL/min. The needle was left in place for 10 min after each injection to allow for diffusion of infusate. For the LEC sites, single fibre optic cannulae (Doric Lenses Inc.) were implanted bilaterally. To allow for sufficient clearance to connect both left and right optic fibres to the light source, the LEC implants were angled away from one another – 30° from the AP-axis – i.e., one pointed forward and one pointed back (this was counterbalanced between hemispheres). Additionally, they were angled 20° to the vertical in the ML axis, to allow access to LEC with minimal damage to perirhinal cortex. Thus, the implantation site began at a different location on each hemisphere; for one hemisphere the burr hole was AP-7.6 mm (connector pointing posteriorly), while on the other it was AP-4.7 mm from bregma (connector pointing anteriorly) (Figure 2A). On both sides the burr hole was ±3.6 mm from the midline and the optic fibre was lowered 8 mm from the surface of the skull along the depth axis of the stereotaxic frame manipulator arm. The optic fibre cannulae were anchored to the skull by stainless steel skull screws (Plastics One) and dental acrylic. After surgery, each animal was given fluid replacement therapy of at least 5 ml of glucose saline subcutaneously and analgesia intramuscularly (0.05 ml of 0.3 mg/ml buprenorphine). Animals were housed individually for 7 days to recover from surgery and in pairs thereafter. The animals recovered for at least 14 days before habituation to the testing arena began.

**Figure 2:**
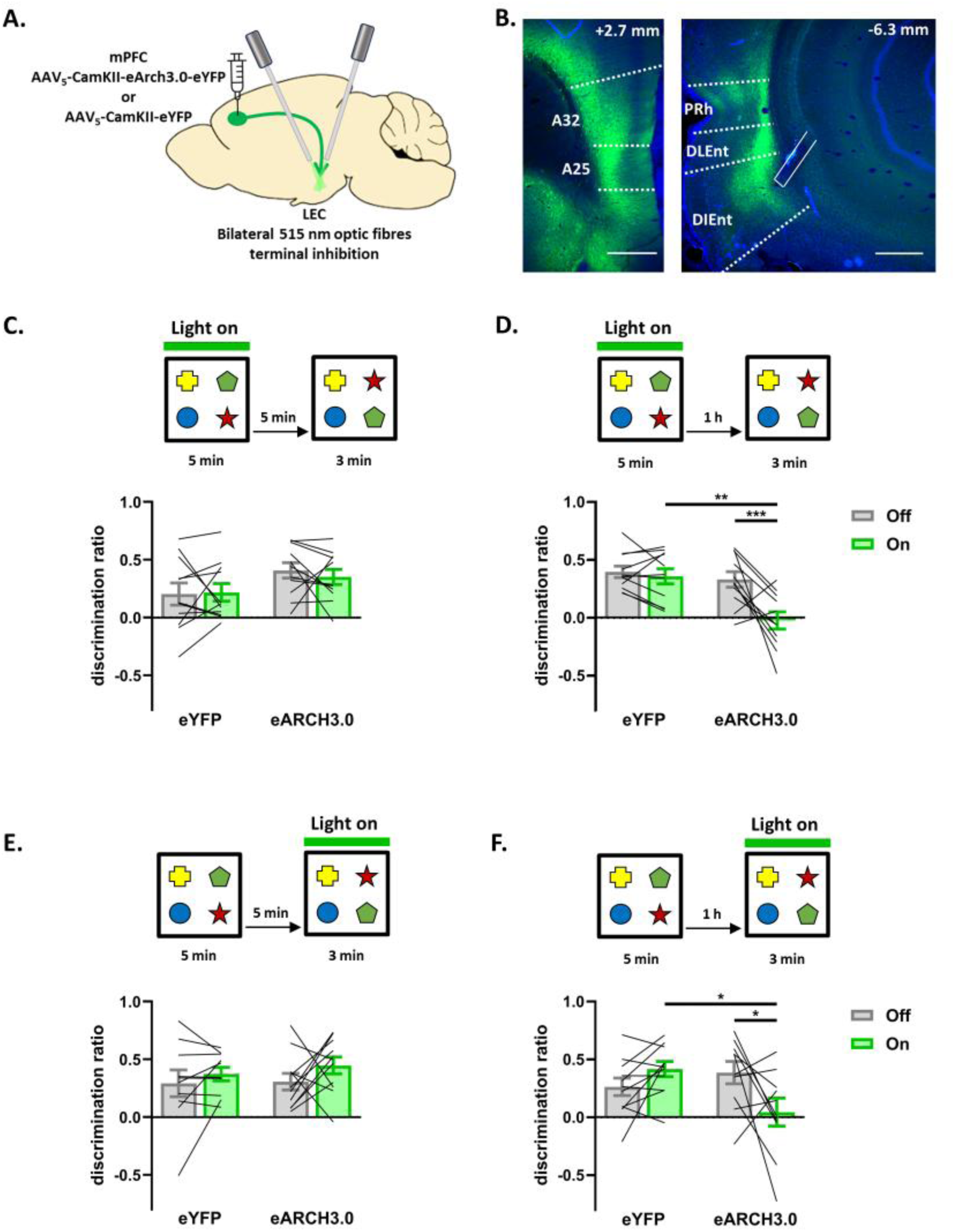
Optogenetic inhibition of mPFC→LEC projections impairs encoding and retrieval of object-in-place in a delay-dependent manner. **A**: Schematic of bilateral mPFC viral injection and LEC fibre-optic implantataion. **B:** Representative image of mPFC viral transduction (left) and YFP positive fibres and angled fibre optic placement in LEC (right). Numbers indicate anteroposterior distance from bregma in mm. Scale = 500 µm. **C:** Optogenetic inhibition of mPFC→LEC during sample does not impair OiP memory with a 5 min delay. **D:** Optogenetic inhibition of mPFC→LEC during sample impairs OiP memory with a 1 h delay. **E:** Optogenetic inhibition of mPFC→LEC during test does not impair OiP memory with a 5 min delay. **F:** Optogenetic inhibition of mPFC→LEC during test impairs OiP memory with a 1 h delay

### Examination of cannulation/optic fibre locations

On completion of the behavioural tasks, the animals with indwelling cannulae were infused with 1 µl Alexa-594 hydrazide (1 mM dissolved in saline, Abcam). Animals were anaesthetised with Euthatal (Rhone Merieux) and underwent transcardial perfusion with phosphate buffer (PB) followed by 4% paraformaldehyde (PFA). The brains were postfixed in 4% PFA for a minimum of 24 h, followed by 48 h in 30% sucrose in PB. Coronal sections (40 µm) were cut on a cryostat and the sections mounted directly onto poly-L-lysine coated slides and coverslipped using DPX mounting medium. Sections were then viewed under an epi-fluorescence microscope and the location of the cannula/optics recorded. To assess the extent of mPFC viral transduction, representative sections along the AP axis between 1.7 and 4.2 mm relative to bregma were selected. LEC cannula placement and eYFP transduced projections from mPFC were assessed between –5.0 and –8.0 mm, relative to bregma and compared with those in the rat brain atlas (Paxinos and Watson, 2018).

### Behavioural apparatus

All behavioural testing took place in a wooden, open-topped arena (50 cm high, 90 cm wide, and 100 cm long) with external black curtains to a height of 1.5 m to restrict distal cues. Object exploration was monitored using an overhead camera connected to a PC and quantified using in-house software on a computer within the room. New objects were used for every experiment.

For the OiP and novel object preference task (NOP) one wall of the arena was black (left with respect to the experimenter) while the other three were painted grey. The floor was covered in sawdust and for the OiP task the curtains were partially removed from around the arena to provide additional extra-maze cues. Objects were constructed from Duplo Lego (Lego^TM^, Denmark) and varied in colour and size from 16 x 16 x 8 cm to 20 x 20 x 25 cm and were too heavy to displace.

To provide different contexts for the OiC task, a smaller behavioural arena was placed within in the arena described above. These arenas measured 50 cm in each dimension; one had black and white vertical striped walls with small white cardboard chips on the floor (‘context A’), the other had black walls with a sawdust floor (‘context B’). Objects were made from Duplo Lego (Lego^TM^, Denmark) and varied in colour and size from 8 x 8 x 5 cm to 11 x 12 x 8 cm. To prevent movement the objects were secured to the floor of the arena using hook and loop tape.

### Behavioural procedures

#### Habituation

Post-surgery all rats were handled daily. Fourteen days following surgery they were habituated to the testing arena without stimuli for 10 min before the commencement of the behavioural testing. The animals were also habituated to the infusion/connection procedure.

For the object-in-context task animals received additional habituation to both contexts across 4 days, each day comprised a 5 min morning and 5 min afternoon session. On day 1, half of the rats received both sessions in context A and the other half saw context B. On day 2, both sessions were in the opposite context to day 1. Days 3 and 4 were mixed sessions, e.g., the morning session in context A and afternoon session in context B (or vice versa) and the opposite on the following day.

#### Object-in-place

The object-in-place (OiP) task comprised a 5min sample phase and 3min test phase separated by a variable delay. In the sample phase each subject was presented with four different objects placed towards the corners of the arena, 10 cm from the walls allowed to explore the objects. During the delay period (5 min or 1 h), objects were cleaned with ethanol to remove olfactory cues and any sawdust that had stuck to the object. In the test phase, two of the objects; for example, objects B and D (which were both on the left or right of the arena), exchanged positions (Figure 1C) and the subject was allowed to explore the objects. The time spent exploring the two objects that had changed position was compared with the time spent exploring the two objects that had remained in the same position. The objects moved (i.e., those on the left or right) and the position of the objects in the sample phase were counterbalanced between rats. If OiP memory is intact, then the subject will spend more time exploring the two displaced objects compared with the two objects that are in the same locations.

#### Novel object preference

Novel object preference (NOP) testing comprised a 5 min sample and 3 min test phase separated by a 1 hour delay. In the sample phase, each subject was presented with four different objects. For the test phase, two of these objects (either the pair on the left or the right) were replaced with novel objects (Figure 1G). The positions of the objects in the test and the objects used as novel or familiar were counterbalanced between the animals. If NOP memory is intact, then animals will spend longer exploring the novel compared with the familiar objects.

#### Object-in-context

The object-in-context (OiC) task comprised of two 4 min sample phases, separated by 5 min followed by a 3 min test phase (Figure 3A). In sample phase 1 animals were placed in one context, e.g., context A. containing two identical copies of an object located near two corners and were allowed to explore the objects for 4 min before being removed from the arena. In sample phase 2, the rats were placed in context B, and allowed to explore two identical copies of a different object for 4 min before being removed from the arena After a 5 min or 1 hour retention delay, the animals were placed back into context A containing one object from sample phase 1 – the congruent object – and one object from sample phase 2 – the incongruent object (Figure 3A). The order of contexts, objects seen within each context and the side of the incongruent object were counterbalanced. If memory for the objects’ original context is intact, then animals should spend longer exploring the incongruent object.

**Figure 3:**
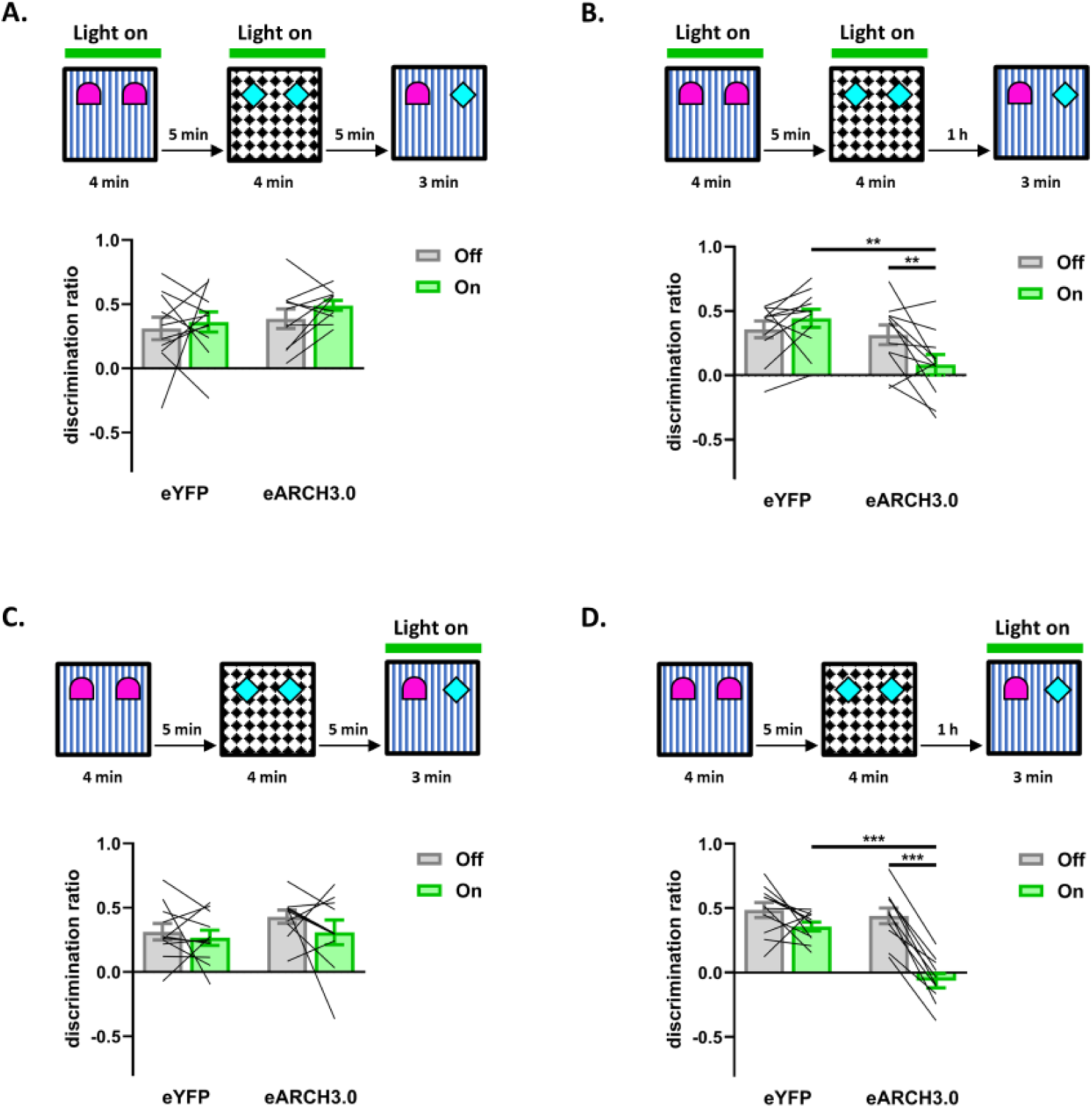
Optogenetic inhibition of mPFC→LEC projections impairs encoding and retrieval of object-in-context in a delay-dependent manner. **A**: Optogenetic inhibition of mPFC→LEC during sample does not impair OiC memory with a 5 min delay **B:** Optogenetic inhibition of mPFC→LEC during sample impairs OiC memory with a 1 h delay **C:** Optogenetic inhibition of mPFC→LEC during test does not impair OiC memory with a 5 min delay **D:** Optogenetic inhibition of mPFC→LEC during test impairs OiC memory with a 1 h delay

#### Two-context novel object preference task

This task comprised two 4 min sample phases and a 3 min test phase separated by a 1 hour delay (Figure 4B). The test phase took place in the context seen for sample phase 1, the object seen in sample phase 1 was presented with a novel object. Object exploration was recorded for 3 min. If memory is intact, the animals will spend more time exploring the novel object.

**Figure 4:**
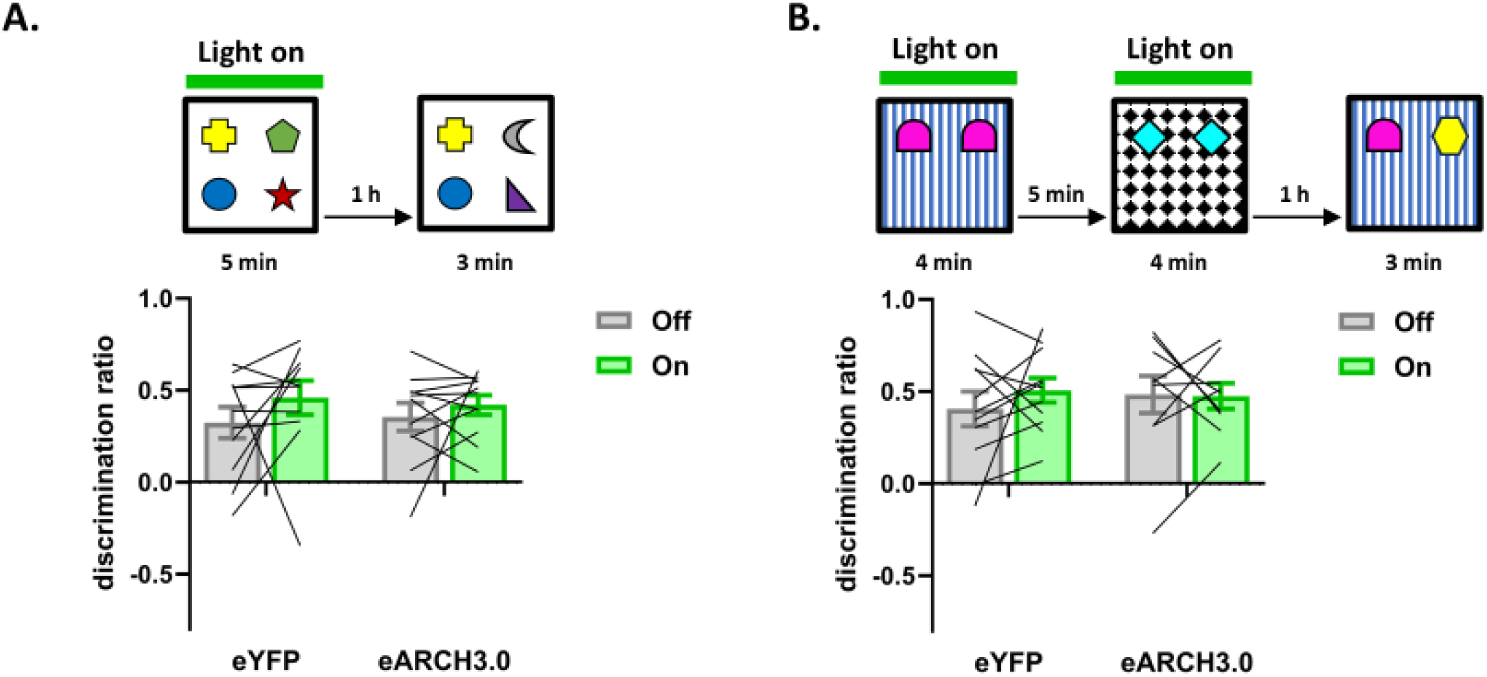
Optogenetic inhibition of mPFC→LEC projections does not affect novel object preference. **A**: Optogenetic inhibition of mPFC→LEC during sample phase of a four-object novel object preference task does not impair memory **B:** Optogenetic inhibition of mPFC→LEC during sample phase of a two-context novel object preference task does not impair memory

#### Drug infusions

General procedures followed those of Barker and Warburton (Barker and Warburton, 2018). The drugs used was 2,3-Dioxo-6-nitro-1,2,3,4-tetrahydrobenzo[f]quinoxaline-7-sulfonamide (NBQX; Biotechne-Tocris) dissolved in sterile 0.9% saline solution. Vehicle infusions consisted of sterile 0.9% saline. NBQX was infused at a concentration of 1 mM. Infusions were made through a 33-gauge cannula (Plastics One) inserted into the implanted guide cannulae and extending 1 mm beyond the end of the guide cannula and attached to a 5 µl Hamilton syringe via polyethylene tubing. A volume of 1 µl of fluid was injected into the mPFC and/or LEC over a 2 min period using an infusion pump (Harvard Bioscience). After the infusion period, the infusion cannula remained in place for a further 5 min before being removed. To test effects of regional temporary inactivation on encoding, the infusions were made 15 min before the sample phase. To determine effects on retrieval, infusions were made 15 min before the test phase (8 min after cannula removal).

#### Optogenetic stimulation protocol

Laser light for optical stimulation was generated using a diode laser (Omicron LuxX® 515-100 laser (515nm), Photonlines, UK). The laser was attached to a fibre optic rotary joint with beam splitter (FRJ 1X2i FC-2FC, Doric Lenses, Quebec, Canada) via a fibre-optic patch cord (core = 200µm, numerical aperture = 0.22, FG200LEA, ThorLabs, Newton, NJ, USA). Two fibre-optic patch cords (core = 200 µm, numerical aperture = 0.22, FC-CM3, Doric Lenses, Quebec, Canada) were attached to the rotary joint at one end while the other end was used to connect to the optical implant on the animal’s head. The power output of the laser was adjusted so that 12 mW was measured at the tip of each optical fibre. Optical stimulation was either given during the sample phase to test the effects on encoding or during the test phase to test the effect on retrieval. Laser stimulation was delivered at a frequency of 50 Hz and a duration of 10 ms to give a 50% duty cycle (Barker et al., 2021) using a custom protocol on WinLTP2.20 software (Anderson and Collingridge, 2007).

### Experimental Design

To assess the individual requirement for mPFC and LEC for successful recognition memory (OiP or NOR), rats received bilateral infusions of either drug or vehicle into each region in separate trials. The control group was made up of animals which received a vehicle infusion into mPFC and animals which received a vehicle infused in LEC. The order of bilateral mPFC infusion, bilateral LEC infusion or vehicle infusion was counterbalanced.

To assess the need for a functional interaction between the mPFC and LEC in the recognition memory tasks, the drugs were infused unilaterally into the mPFC and LEC in either the same hemisphere (IPSI) or in opposite hemispheres (CONTRA). All experiments were run using a cross-over design and each animal re-tested following a rest period of a week. The initial group size was 12 animals. Cannula blockage or misplaced cannulae, following histological verification, resulted in the occasional loss of animals from the study, details of animal exclusions are provided in the results. To assess whether the projection from the mPFC to the LEC was necessary for recognition memory, optogenetic experiments were run with a cross-over design. Half the animals received the virus with the opsin (AAV5-CaMKii-eARCH3.0-eYFP) and half received the control virus (AAV5-CaMKii-eYFP). All animals were tested with both optical stimulation-on and stimulation-off conditions in a cross-over design.

### Analysis and Statistics

The duration of the exploratory behaviour (in seconds) of the objects was defined as the animal directing its nose toward the object at <2 cm from the object. Other behaviours, such as sitting on or resting against the object or using it for supported rearing were not included. Animals were excluded from analysis of a particular task if they failed to explore all the objects or failed to reach a minimum total exploration time of 15 seconds in the sample phase(s) and 10 seconds in the test phase. Measurement of object exploration in the test phase was used to calculate a discrimination ratio comparing exploration of **novel** (i.e. novel object /novel object-place configuration/ object in a novel context (incongruent object)) with **familiar** (i.e. familiar object/familiar object-place configuration/ object in the familiar context (congruent object)). This ratio was calculated using the formula: novel exploration – familiar exploration/ total exploration time (novel+familiar). This ratio was chosen as it takes individual differences in the total object exploration into account. A value of zero indicates that the animal has no preference for the novel or familiar, while a positive discrimination ratio value indicates that an animal has preference for the novel.

Statistical comparisons between group exploration times and group discrimination ratios were made using paired Student’s t-tests (two-tailed) or analysis of variance (ANOVA; SPSS version 24, IBM) as appropriate based on experimental design. In the pharmacological experiments, the within-subjects factor was treatment (mPFC/LEC/vehicle) or group (IPSI vs. CONTRA). If a significant effect of treatment was found post-hoc comparisons used a Holm correction for multiple comparisons. In the optogenetic experiments the within-subjects factor was laser stimulation condition (ON vs. OFF) and the between-subjects factor was virus (eARCH3.0 vs. eYPF). We also examined the total object exploration levels in the sample and test phases of each recognition memory test. To examine the whether the amount of object exploration in the sample phase may have impacted the results in the OiC experiments, a three-way ANOVA was used with stimulation, virus and sample phase as factors.

When the interaction term was found to be significant, post hoc simple main effects tests were calculated. Whether individual groups discriminated between the objects was assessed using a one-sample Student’s t tests (two-tailed) to determine if the discrimination ratio differed significantly from zero (chance level). All statistical analyses used a significance level of 0.05.

## Results

### Experiment 1: pharmacological inactivation of mPFC and LEC

#### Histological verification of cannula placements

Histological examination confirmed that all cannulae implanted in the mPFC had needle tips located in the ventral portion of prelimbic and dorsal portion of the infralimbic regions (Figure 1B). However, in two cases, the cannulae aimed at the LEC were located beyond the LEC border and so the data from those rats were excluded from all analyses, therefore n = 10 for this cohort.

### mPFC and LEC are individually required for both encoding and retrieval of object-in-place memory

Bilateral administration of the AMPA receptor antagonist NBQX into either the mPFC or LEC before sample, to temporarily inactivate these regions during memory encoding, significantly impaired OiP performance following a 1h delay between sample and test (one-way ANOVA F_2,18_ = 4.33, p = 0.029; Figure 1C). Post-hoc tests revealed that NBQX infusion into the LEC produced a significant impairment in performance compared to vehicle (p = 0.029). However, there was no significant difference following infusions into the mPFC (p = 0.16). Further analyses compared the group discrimination ratios to chance levels, to check whether each group showed a significant preference for the novel compared to the familiar object-place configurations. The control group, as predicted showed a clear preference for the novel object-place configuration (t_9_ = 3.91, p = 0.0035) while the discrimination ratios of the mPFC and LEC groups was not significantly different from zero indicating that neither group significantly discriminated (mPFC: t_9_ = 1.04, p = 0.33; LEC: t_9_ = –0.27, p = 0.80).

Administration of NBQX, into the mPFC or LEC, prior to the test phase similarly impaired OiP performance (F_2,18_ = 4.85, p = 0.021; Figure 1D). Post-hoc tests confirmed the significant impairment following infusions of NBQX into the mPFC (p = 0.040) and LEC (p = 0.034), compared to vehicle. As expected, the performance of the control group was significantly above chance (t_9_ = 2.64, p = 0.027) while that of the mPFC and LEC groups was not (mPFC: t_9_ = 0.029, p = 0.98; LEC: t_9_ = –0.47, p = 0.65).

#### Total Object Exploration

We examined whether administration of NBQX into either the mPFC or LEC, at any stage of the procedure, impacted overall object exploration levels. One-way ANOVA revealed that pre-sample administration of NBQX had no effect on object exploration in the sample (F_2,18_ = 0.27, p = 0.77) or test phases (F_2,18_ = 1.99, p = 0.17; Table 1). Similarly, there was also no effect of pre-test infusions on sample (F_2,18_ = 0.69, p = 0.52) or test phase exploration (F_2,18_ = 0.32, p = 0.73; Table 1).

**Table 1.** Object exploration in the object-in-place task bilateral with infusions of NBQX or vehicle.

| Infusion timing | Site of infusion | Encoding session Exploration (s) | SEM | Test session Exploration (s) | SEM |
| --- | --- | --- | --- | --- | --- |
| Encoding | LEC | 76.26 | ±7.99 | 39.16 | ±4.59 |
|  | mPFC | 81.38 | ±7.11 | 30.48 | ±3.66 |
|  | Vehicle | 74.28 | ±6.60 | 29.72 | ±2.27 |
| Retrieval | LEC | 61.94 | ±4.41 | 30.77 | ±4.19 |
|  | mPFC | 67.98 | ±5.19 | 27.30 | ±3.76 |
|  | Vehicle | 66.49 | ±4.65 | 29.17 | ±4.01 |

These data confirm that mPFC and LEC are each required for both encoding and retrieval of OiP memory.

### The mPFC-LEC circuit is required for both encoding and retrieval of object-in-place memory

To address the question of whether successful OiP memory requires a functional interaction between the mPFC and LEC we next used a disconnection procedure. In these experiments NBQX was infused unilaterally into both the mPFC and LEC in either the same (IPSI) or opposite (CONTRA) hemispheres, so that in the latter group, transmission in a mPFC-LEC circuit across both hemispheres is disrupted. The IPSI group serve as the control group. If the mPFC-LEC are functionally interdependent during OiP, then the performance of the CONTRA group should be impaired compared to the IPSI group (Barker et al., 2007; Barker and Warburton, 2011).

Both pre-sample and pre-test administration of NBQX significantly impaired performance in the CONTRA compared to the IPSI group (pre-sample t_9_ = 2.40, p = 0.040; Figure 1E; pre-test t_9_ = 3.57, p = 0.0060; Figure 1F). Further analysis comparing the discrimination ratios to zero, showed that, pre-sample and pre-test infusions resulted in the CONTRA group showing no preference for the novel compared to the familiar object-place configurations (pre-sample: t_9_ = –0.18, p = 0.86; pre-test: t_9_ = –1.67, p = 0.13), while discrimination in the IPSI group was significantly above chance (pre-sample: t_9_ = 4.84, p = 0.00092; pre-test: t_9_ = 4.16, p = 0.0024).

#### Total Object Exploration

Examination of drug effects on overall object exploration, revealed that pre-sample administration of NBQX resulted in a difference in the sample phase between the IPSI and CONTRA groups which just missed significance (t_9_ = –2.23, p = 0.053). Observation of the group means reveals that, in this case, the CONTRA group explored the objects more than the IPSI group (see Table 2). There was no difference in exploration levels in the test phase (t_9_ = 0.077, p = 0.94; Table 2). Pre-test NBQX administration also did not affect total object exploration in the sample (t_9_ = 0.84, p = 0.42) or test phases (t_9_ = 0.46, p = 0.66; Table 2).

**Table 2.** Object exploration in the object-in-place and novel object preference tasks with contralateral or ipsilateral mPFC-LEC NBQX infusions.

| Task | Infusion | Infusion timing | Site of infusion | Encoding session Exploration (s) | SEM | Test session Exploration (s) | SEM |
| --- | --- | --- | --- | --- | --- | --- | --- |
| OiP | NBQX | Encoding | Contra | 55.44 | ±3.58 | 23.67 | ±3.22 |
|  |  |  | Ipsi | 48.80 | ±3.49 | 24.04 | ±3.56 |
|  |  | Retrieval | Contra | 54.81 | ±4.40 | 32.34 | ±3.69 |
|  |  |  | Ipsi | 58.12 | ±4.72 | 34.67 | ±5.57 |
| NOP | NBQX | Encoding | Contra | 50.57 | ±5.92 | 25.20 | ±2.94 |
|  |  |  | Ipsi | 50.57 | ±6.41 | 30.67 | ±1.68 |

Together these data demonstrate that communication between mPFC and LEC is required for both the encoding and for the retrieval of OiP memory.

### The mPFC-LEC circuit is not required for novel object recognition memory

We next examined whether a mPFC-LEC interaction is necessary for encoding of novel object recognition memory. Pre-sample administration of NBQX in the CONTRA or IPSI groups had no significant effect on novel object recognition memory (t_7_ = 0.42, p = 0.69; Figure 1G); and as both the CONTRA and IPSI groups had discrimination ratios that were significantly above chance (IPSI t_7_ = 5.36, p = 0.0011; CONTRA t_7_ = 4.90, p = 0.0018) both groups were clearly able to discriminate the novel from the familiar objects.

#### Total Object Exploration

Pre-sample infusion of NBQX did not alter total group object exploration times in either the sample (t_7_ = 0.038, p = 0.97) or test phases (t_7_ = 2.01, p = 0.085; Table 2).

Thus, communication between mPFC and LEC is not required for novel object recognition memory.

### Experiment 2: The role of the mPFC axonal terminals in the LEC in short and long-term object-in-place memory

Disconnection analysis showed the importance of the mPFC-LEC interaction for associative recognition memory, however it did not reveal the direction of information transfer between these regions. Hence, we next used optogenetic inactivation to test the necessity of the direct mPFC→LEC pathway given that mPFC efferents to the nucleus reuniens or medial dorsal thalamus are selectively required for OiP retrieval (Barker et al., 2021; Culleton et al., unpublished). The temporal control afforded by the optogenetic technique also allowed us to examine memory performance following either a 5min or a 1h delay.

### Histological verification of viral injection and optical fibre placements

On completion of the behavioural experiments, the extent of eYFP expression and optical fibre placement were examined. In all animals, injection of AAV-ARCH-eYFP or AAV-eYFP resulted in robust expression in the mPFC with the strongest expression seen in layers 5 and 6 of A25 and A32 (Figure 2A and 2B). Axonal transport of the eYFP reporter was observed in the LEC, where mPFC efferents were primarily located in layers 5 and 6, in all animals (Figure 2B). In two animals (one from the eARCH and one from the eYFP group) the optical fibres were located outside of the LEC, and these animals were excluded from statistical analysis, therefore n = 11 rats in both the eYFP and eARCH groups. All other animals had the optical fibres located to enable inhibition of mPFC terminals in the LEC. Based on the optogenetic parameters, it is predicted that a minimum of 10 mWmm^-2^ light density would be produced within a 0.5 mm range from the tip of the optical fibre (Gradinaru et al., 2007; Deisseroth, 2012) (Deisseroth, 2012; Gradinaru et al., 2009) which is capable of robustly activating eARCH3.0 (Mattis et al., 2011).

### Optogenetic inactivation of PFC-LEC transmission during the encoding phase of the object-in-place task

Optogenetic inhibition of mPFC axon terminals in the LEC during the encoding phase of the OiP task impaired memory in a delay dependent manner (Figure 2C, 2D). Statistical analysis of group discrimination ratios revealed that light delivery during the sample phase, had no effect on performance following a 5min delay (condition x virus interaction (F_1,20_ = 0.34, p = 0.57), no significant main effect of stimulation (F_1,20_ = 0.12, p = 0.73) or virus (F_1,20_ = 3.50, p = 0.076). Further, all groups successfully discriminated between the novel and familiar object-place configurations (eYFP-OFF: t_10_ = 2.54, p = 0.027; eYFP-ON: t_10_ = 3.26, p = 0.0075; eARCH3.0-OFF: t_10_ = 6.12, p < 0.00011; eARCH3.0-ON: t_10_ = 5.44, p < 0.00029).

Following the 1-hour retention delay, analysis of the discrimination ratios found a significant stimulation x virus interaction (F_1,20_ = 7.06, p = 0.015; Figure 2D) and significant main effect of stimulation (F_1,20_ = 11.03, p =0.0034) and virus (F_1,20_ = 10.25, p = 0.0045). Analysis of the simple main effects revealed that this interaction was driven by a significantly lower discrimination ratio in the eARCH3.0-ON group when compared to both the eARCH3.0-OFFgroup (p = 0.00041) and the eYFP-ON group (p = 0.0011). Analysis comparing performance against chance revealed that all groups except for the eARCH3.0-ON group successfully discriminated between the novel and familiar object-place configurations (eYFP-OFF: t_10_ = 8.07, p = 0.000011; eYFP-ON: t_10_ = 5.36, p = 0.00032; eARCH3.0-Off: t_10_ = 4.92, p = 0.001; eARCH3.0-On: t_10_ = 0.35, p = 0.74).

Together these results indicate that mPFC input to the LEC is required for the encoding of OiP information retained after a 1h but not 5 min delay.

#### Total Object Exploration

<u>Five minute delay</u> – Analysis of the overall amount of object exploration completed during the sample phase revealed no significant interaction between stimulation and virus (F_1,20_ = 0.51, p = 0.48) and no significant main effect of stimulation (F_1,20_ = 1.08, p = 0.31) or virus (F_1,20_ = 0.51, p = 0.48; Table 3). Analysis of the amount of object exploration completed in the test phase showed there was no significant interaction between stimulation and virus (F_1,20_ = 0.04, p = 0.84) and no significant main effect of stimulation (F_1,20_ = 0.45, p = 0.51) or virus (F_1,20_ = 4.14, p = 0.055; Table 3).

**Table 3.** mPFC→LEC terminal inhibition experiment object exploration, object-in-place and four-object novel object preference tasks.

| Task | Stimulation |  | Virus | Laser | Encoding session |  | Test session |  |
| --- | --- | --- | --- | --- | --- | --- | --- | --- |
|  | timing | Delay |  |  | Exploration (s) | SEM | Exploration (s) | SEM |
| OiP | Encoding | 1 hour | eArch3.0 | Off | 75.11 | ±4.99 | 32.89 | ±4.15 |
|  |  |  |  | On | 78.05 | ±5.67 | 33.81 | ±3.37 |
|  |  |  | eYFP | Off | 73.98 | ±4.01 | 33.49 | ±2.36 |
|  |  |  |  | On | 74.57 | ±6.02 | 28.99 | ±3.14 |
|  |  | 5 mins | eArch3.0 | Off | 78.85 | ±3.47 | 50.77 | ±5.90 |
|  |  |  |  | On | 72.91 | ±5.79 | 47.32 | ±3.74 |
|  |  |  | eYFP | Off | 74.47 | ±6.69 | 40.84 | ±4.14 |
|  |  |  |  | On | 73.38 | ±4.92 | 38.99 | ±2.41 |
|  | Retrieval | 1 hour | eArch3.0 | Off | 77.11 | ±4.32 | 36.38 | ±2.46 |
|  |  |  |  | On | 69.08 | ±3.71 | 29.98 | ±3.86 |
|  |  |  | eYFP | Off | 79.90 | ±5.29 | 26.87 | ±2.49 |
|  |  |  |  | On | 70.23 | ±5.85 | 31.24 | ±2.09 |
|  |  | 5 mins | eArch3.0 | Off | 91.96 | ±6.82 | 42.59 | ±3.90 |
|  |  |  |  | On | 85.65 | ±4.48 | 44.73 | ±3.68 |
|  |  |  | eYFP | Off | 94.29 | ±7.16 | 48.24 | ±4.98 |
|  |  |  |  | On | 89.71 | ±6.48 | 46.02 | ±4.14 |
| NOP | Encoding | 1 hour | eArch3.0 | Off | 83.42 | ±5.81 | 47.30 | ±4.38 |
|  |  |  |  | On | 87.46 | ±5.83 | 43.92 | ±3.27 |
|  |  |  | eYFP | Off | 99.40 | ±4.59 | 48.83 | ±3.71 |
|  |  |  |  | On | 94.06 | ±6.31 | 48.52 | ±4.72 |

<u>One hour delay</u>-Analysis of the overall amount of object exploration completed during the sample phase revealed no significant interaction between stimulation and virus (F_1,20_ = 0.13, p = 0.72) and no significant main effect of stimulation (F_1,20_ = 0.30, p = 0.59) or virus (F_1,20_ = 0.13, p = 0.72; Table 3). In addition, analysis of object exploration in the test phase also revealed no significant interaction between stimulation and virus (F_1,20_=0.77, p=0.39), no significant main effect of stimulation (F_1,20_ = 0.34, p = 0.57) or virus (F_1,20_ = 0.77, p = 0.39; Table 3).

### Optogenetic inactivation of PFC-LEC transmission during the retrieval phase of the object-in-place task

To examine the effects of inhibition of the mPFC input to the LEC during the retrieval phase of the OiP task (Figure 2E, 2F) light was delivered during the test phase, 5 min or 1 h after the sample phase. When there was a 5 min delay between the sample and test phases discrimination performance was not impaired, as confirmed by statistical analysis showing no significant stimulation x virus interaction (F_1,19_ = 0.16, p = 0.69; Figure 2E), no significant main effect of stimulation (F_1,19_ = 2.23, p = 0.15) or virus (F_1,19_ = 0.24, p = 0.63). Further analyses showed that all groups successfully discriminated between the novel and familiar object-place configurations (eYFP-Off: t_9_ = 2.52, p = 0.033; eYFP-On: t_9_ = 6.40, p = 0.00013; eARCH3.0-Off: t_10_ = 4.27, p = 0.0016; eARCH3.0-On: t_10_ = 6.17, p = 0.00011). When there was a 1h delay between the sample and test, light delivery produced a significant impairment in performance. Statistical analysis revealed a significant stimulation x virus interaction (F_1,19_ = 7.51, p = 0.013; Figure 2F) but no significant main effect of stimulation (F_1,19_ = 1.10, p = 0.31) or virus (F_1,19_ = 1.90, p = 0.18). Simple main effects analysis revealed this was due to a significantly lower discrimination ratio in the eARCH3.0-ON group than both the eARCH3.0-OFF group (p = 0.017) and the eYFP-ON group (p = 0.012). Further analysis comparing performance against chance again showed that all control groups successfully discriminated above chance level (eYFP-OFF: t_10_ = 3.47, p = 0.006; eYFP-ON: t_10_ = 6.35, p = 0.000083; eARCH3.0-OFF: t_9_ = 4.00, p = 0.0016). However, when mPFC terminals in LEC were inhibited in the eARCH3.0-On group the discrimination ratio was no different from chance (eARCH3.0-ON: t_9_ = 0.35, p = 0.73). Thus, mPFC input to the LEC during memory retrieval is required for successful OiP memory only following a 1h delay.

#### Total Object Exploration

<u>Five minute delay</u> –Analysis of total object exploration completed during the sample phase revealed no significant stimulation x virus interaction (F_1,19_ = 0.032, p = 0.86; Table 3) and no significant main effect of stimulation (F_1,19_ = 1.27, p = 0.27) or virus (F_1,19_ = 0.18, p = 0.67). Similarly, analysis of total object exploration during the test phase found no significant stimulation x virus interaction (F_1,19_ = 0.34, p = 0.57) or main effects of stimulation (F_1,19_ = 0.000086, p = 0.99) or virus (F_1,19_ = 0.58, p = 0.46) (Table 3).

<u>One hour delay</u> – Analysis of the overall amount of object exploration completed during the sample phase revealed no significant interaction between stimulation x virus (F_1,19_ = 0.091, p = 0.77; Table 3) and no significant main effect of virus (F_1,19_ = 0.093, p = 0.76). There was a significant main effect of stimulation (F_1,19_ = 10.62, p = 0.0041) which reflected decreased levels of exploration in both eYFP and eARCH3.0 groups when the laser was on. Analysis of the total amount of object exploration during the test phase revealed a significant stimulation x virus interaction (F_1,19_ = 6.07, p = 0.023; Table 3) but no significant main effect of stimulation (F_1,19_ = 0.22, p = 0.65) or virus (F_1,19_ = 1.62, p = 0.22). Examination of the simple main effects showed this significant interaction was driven by a significant difference between the eARCH3.0 and eYPF groups in the light-OFF condition (p = 0.014) with the eARCH3.0 group completing a greater amount of object exploration compared to the eYFP group. Importantly there was no significant difference in the amount of object exploration completed in the test phase of animals expressing eARCH3.0 when the laser was off or on (p = 0.057).

### Optogenetic inactivation of PFC-LEC transmission during object-in-context memory

One animal in the eARCH group was excluded due to the loss of an optical fibre, hence there was a reduced group size for the OiC 5 min sample phase stimulation experiment, both OiC test phase stimulation experiments and the two-context object recognition experiments.

### Optogenetic inactivation of PFC-LEC transmission during the encoding phases of the object-in-context task

To explore whether the delay-dependent requirement of the mPFC-LEC pathway in OiP memory may be generalised to other forms of associative recognition memory, we next tested the same animals in a series of OiC recognition memory tasks. Here animals encountered objects within specific contexts during two sample phases. At test the amount of time spent exploring the incongruent and congruent objects was compared. Light stimulation was delivered either during the sample phases or during the test phase.

Analysis of the discrimination ratios following the 5 min delay between encoding and retrieval revealed a non-significant stimulation x virus interaction (F_1,19_ = 0.13, p = 0.72; Figure 3A) main effect of stimulation (F_1,19_ = 1.15, p = 0.30) and virus (F_1,19_ = 1.82, p = 0.19). In addition, all groups successfully discriminated between the congruent and incongruent objects (eYFP-OFF: t_10_ = 3.57, p = 0.0051; eYFP-ON: t_10_ = 4.61, p = 0.0010; eARCH3.0-OFF: t_9_ = 5.05, p = 0.00069; eARCH3.0-ON: t_9_ = 12.7, p = 4.8 x10^-7^). Analysis of the discrimination ratios following a 1-hour retention delay revealed a significant stimulation condition x virus interaction (F_1,20_ = 8.93, p = 0.007; Figure 3B) and significant main effect of virus (F_1,20_ = 5.22, p = 0.033) but no significant main effect of stimulation condition (F_1,20_ = 1.90, p = 0.18). Examination of the simple main effects revealed that this interaction was driven by a significantly lower discrimination ratio in the eARCH3.0-ON group when compared to both the eARCH3.0-OFF group (p = 0.0058) and the eYFP-ON group (p = 0.0028). Analysis comparing performance against chance revealed that all the control groups successfully discriminated between the incongruent and congruent objects (eYFP-OFF: t_10_ = 5.46, p = 0.00028; eYFP-ON: t_10_ = 6.27, p = 0.000093; eARCH3.0-OFF: t_10_ = 4.13, p = 0.0020). However, the eARCH3.0-ON group did not discriminate above chance level (t_10_ = 1.02, p = 0.33) indicative of a significant memory impairment in this group.

#### Total object exploration

<u>Five minute delay</u>-There was no significant stimulation x virus x sample phase interaction (F_1,19_ = 0.056, p = 0.82), stimulation x virus interaction (F_1,19_ = 0.057, p = 0.81), stimulation x sample phase (F_1,19_ = 0.014, p = 0.91); virus x sample phase F_1,19_ = 0.097, p = 0.78) and no significant main effect of stimulation (F_1,19_ = 3.69, p = 0.070), virus (F_1,19_ = 0.21, p = 0.65) or sample phase (F_1,19_ = 0.082, p = 0.78). We also examined object exploration in the test phase and found no significant interaction (stimulation x virus F_1,19_ = 0.0011, p = 0.97) and no significant main effects (stimulation F_1,19_ = 0.036, p = 0.85; virus F_1,19_ = 0.00084, p = 0.98; Table 4).

**Table 4.**
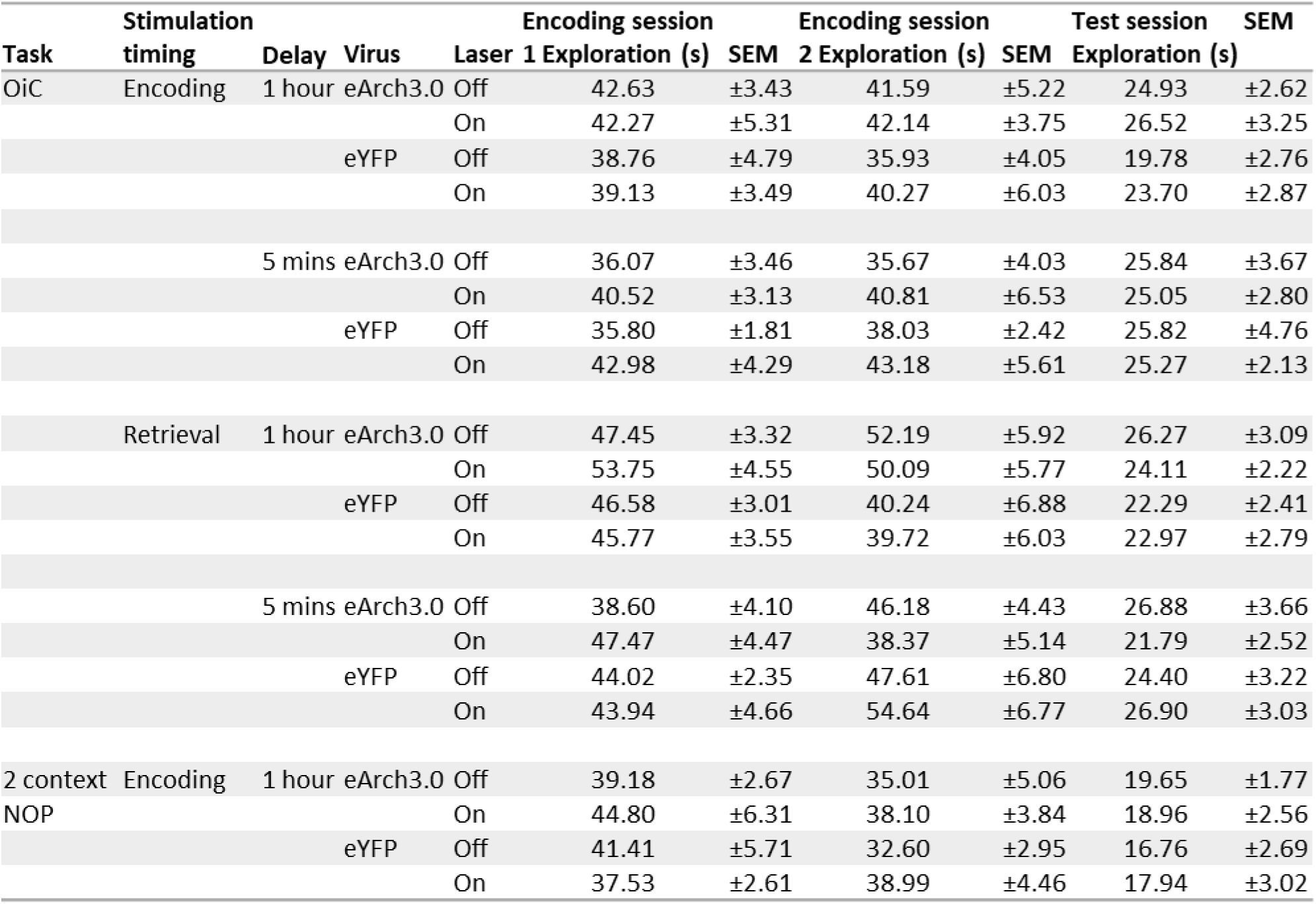
mPFC→LEC terminal inhibition experiment object exploration, object-in-context and two-context novel object preference tasks.

| Task | Stimulation |  | Virus | Laser | Encoding session 1 |  | Encoding session 2 |  | Test session |  |
| --- | --- | --- | --- | --- | --- | --- | --- | --- | --- | --- |
|  | timing | Delay |  |  | Exploration (s) | SEM | Exploration (s) | SEM | Exploration (s) | SEM |
| OiC | Encoding | 1 hour | eArch3.0 | Off | 42.63 | ±3.43 | 41.59 | ±5.22 | 24.93 | ±2.62 |
|  |  |  |  | On | 42.27 | ±5.31 | 42.14 | ±3.75 | 26.52 | ±3.25 |
|  |  |  | eYFP | Off | 38.76 | ±4.79 | 35.93 | ±4.05 | 19.78 | ±2.76 |
|  |  |  |  | On | 39.13 | ±3.49 | 40.27 | ±6.03 | 23.70 | ±2.87 |
|  |  | 5 mins | eArch3.0 | Off | 36.07 | ±3.46 | 35.67 | ±4.03 | 25.84 | ±3.67 |
|  |  |  |  | On | 40.52 | ±3.13 | 40.81 | ±6.53 | 25.05 | ±2.80 |
|  |  |  | eYFP | Off | 35.80 | ±1.81 | 38.03 | ±2.42 | 25.82 | ±4.76 |
|  |  |  |  | On | 42.98 | ±4.29 | 43.18 | ±5.61 | 25.27 | ±2.13 |
|  | Retrieval | 1 hour | eArch3.0 | Off | 47.45 | ±3.32 | 52.19 | ±5.92 | 26.27 | ±3.09 |
|  |  |  |  | On | 53.75 | ±4.55 | 50.09 | ±5.77 | 24.11 | ±2.22 |
|  |  |  | eYFP | Off | 46.58 | ±3.01 | 40.24 | ±6.88 | 22.29 | ±2.41 |
|  |  |  |  | On | 45.77 | ±3.55 | 39.72 | ±6.03 | 22.97 | ±2.79 |
|  |  | 5 mins | eArch3.0 | Off | 38.60 | ±4.10 | 46.18 | ±4.43 | 26.88 | ±3.66 |
|  |  |  |  | On | 47.47 | ±4.47 | 38.37 | ±5.14 | 21.79 | ±2.52 |
|  |  |  | eYFP | Off | 44.02 | ±2.35 | 47.61 | ±6.80 | 24.40 | ±3.22 |
|  |  |  |  | On | 43.94 | ±4.66 | 54.64 | ±6.77 | 26.90 | ±3.03 |
| 2 context NOP | Encoding | 1 hour | eArch3.0 | Off | 39.18 | ±2.67 | 35.01 | ±5.06 | 19.65 | ±1.77 |
|  |  |  |  | On | 44.80 | ±6.31 | 38.10 | ±3.84 | 18.96 | ±2.56 |
|  |  |  | eYFP | Off | 41.41 | ±5.71 | 32.60 | ±2.95 | 16.76 | ±2.69 |
|  |  |  |  | On | 37.53 | ±2.61 | 38.99 | ±4.46 | 17.94 | ±3.02 |

<u>One hour delay</u> – There were no significant interactions between stimulation, virus and sample phase (stimulation x virus x sample phase F_1,20_ = 0.075, p = 0.79; stimulation x virus F_1,20_ = 0.16, p = 0.70; stimulation x sample phase F_1,20_ = 0.19, p=0.67; virus x sample phase F_1,20_ = 0.0038, p = 0.95) and no significant main effect of stimulation condition (F_1,20_ = 0.18, p = 0.67), virus (F_1,20_ = 0.60, p = 0.45) and sample phase (F_1,20_ = 0.12, p = 0.74; Table 4). In addition, no significant interaction (stimulation condition by virus (F_1,20_ = 0.34, p = 0.57) and no significant main effects (stimulation (F_1,20_ = 1.91, p=0.18; virus (F_1,20_ = 1.25, p = 0.28) were found in total object exploration completed in the test phase (Table 4).

Together these results show that mPFC input to the LEC is required for the encoding of OiC memory in a delay dependent manner, being necessary for memory with a 1 h, but not a 5 min, delay.

### Optogenetic inactivation of mPFC→LEC transmission during the retrieval phase of the object-in-place task

One animal (eYFP) was excluded for failing to explore all the objects in the test phase. Analysis of discrimination ratios following a 5min delay revealed no significant stimulation x virus interaction (F_1,19_ = 0.30, p = 0.59; Figure 3C) and no significant main effect of stimulation (F_1,19_ = 1.54, p=0.23) or virus (F_1,19_ = 1.26, p = 0.28). Further, all groups successfully discriminated above chance level (eYFP-OFF: t_10_ = 4.84, p = 0.00069; eYFP-ON: t_10_ = 4.48, p = 0.0012; eARCH3.0-OFF: t_9_ = 8.28, p = 0.000017; eARCH3.0-ON: t_9_ = 3.21, p = 0.011).

One animal (eARCH) was excluded for failing to explore all the objects in the test phase Analysis of discrimination ratios following a 1h delay revealed a significant stimulation x virus interaction (F_1,19_ = 18.8, p = 0.00035; Figure 3D) and a significant main effect of stimulation (F_1,19_ = 52.95, p = 6.7 x10^-7^) and virus (F_1,19_ = 13.44, p = 0.0016). Simple main effects analysis revealed this interaction was due to a significantly lower discrimination ratio in the eARCH3.0-On group than both the eARCH3.0-Off group (p = 1.6 x10^-7^) and the eYFP-ON group (p = 0.000003). Further analysis comparing performance against chance showed that all groups except eARCH3.0-ON successfully discriminated above chance level (eYFP-OFF: t_10_ = 8.40, p = 0.000008; eYFP-ON: t_10_ = 10.3, p = 0.000001; eARCH3.0-OFF: t_9_ = 7.34, p = 0.000025; eARCH3.0-ON: t_9_ = 1.13, p = 0.29).

#### Total exploration

<u>Five minute delay</u> –Analysis of object exploration completed during the sample phase revealed a significant interaction between stimulation x virus x sample phase (F_1,19_ = 0.53 p= 0.032; Table 4) but no significant stimulation x virus interaction (F_1,19_ = 0.44, p= 0.52), stimulation x sample phase (F_1,19_ = 0.86, p = 0.37) and virus x sample phase (F_1,19_ = 1.53, p=0.23). Further there was no significant main effect of stimulation (F_1,19_ = 0.81, p =0.38), virus (F_1,19_ = 0.80, p = 0.38) or sample phase (F_1,19_ = 1.00, p = 0.33). Finally, analysis of exploration during test found no significant stimulation x virus interaction (F_1,19_ = 2.18, p = 0.16), main effect of stimulation (F_1,19_ = 0.25, p = 0.62) or virus (F_1,19_= 0.13, p= 0.72;Table 4).

<u>One hour delay</u>-Analysis of object exploration completed during the sample phases revealed no significant interactions between stimulation x virus x sample phase (F_1,19_ = 0.22, p =0.65); stimulation x virus (F_1,19_ = 0.94, p= 0.35); stimulation x sample phase (F_1,19_ = 0.17, p = 0.69); virus x sample phase (F_1,19_ = 0.94, p = 0.35) and no significant main effect of stimulation (F_1,19_ = 0.54, p =0.47), virus (F_1,19_ = 1.65, p= 0.21) or sample phase (F_1,19_ = 1.79, p = 0.20). Table 4). Analysis of object exploration during the test phase revealed a non-significant x virus interaction (F_1,19_ = 0.091, p = 0.77; Table 4), and non-significant main effects of stimulation (F_1,19_ = 0.00092, p = 0.98) and virus (F_1,19_ = 0.33, p = 0.57).

Together these results indicate that mPFC input to the LEC is required for the retrieval of OiC memory in a delay-dependent manner, i.e. retrieval following a longer but not short delays.

### Optogenetic inactivation of PFC-LEC transmission during the novel object preference task

Finally, we examined the effects of optogenetic inhibition of mPFC axon terminals in the LEC during the encoding phase of two different novel object preference tasks, the method of one version was designed to parallel the OiP task i.e. by using four objects (Figure 4A) and the method of the other version was designed to parallel the OiC task, i.e. by including two-contexts (Figure 4B).

<u>Novel object preference (Four object version)</u> – There was no significant stimulation x virus interaction (F_1,20_ = 0.19, p = 0.66), main effect of stimulation (F_1,20_ = 1.59, p = 0.22) or virus (F_1,20_ = 0.033, p = 0.96; Figure 4A). Further analysis confirmed that all groups successfully discriminated between the novel and familiar objects configurations (eYFP-OFF: t_10_ = 3.83, p = 0.003; eYFP-ON: t_10_ = 4.94, p = 0.00059; eARCH3.0-OFF: t_10_ = 4.65, p = 0.00091; eARCH3.0-ON: t_10_ = 7.86, p = 0.000014).

There were no significant differences in total object exploration in either the sample (stimulation x virus interaction F_1,20_ = 1.05, p = 0.32; stimulation, F_1,20_ = 0.020, p = 0.89; virus F_1,20_ = 2.94, p = 0.10) or test phase (interaction F_1,20_ = 0.20, p = 0.66; stimulation F_1,20_ = 0.28, p = 0.60; virus F_1,20_ = 0.45, p = 0.51; Table 3).

<u>Novel object preference (Two-context version)</u> – One eARCH3.0 animal was removed from the analysis for failing to complete 10 s of object exploration in the test phase. There was no significant stimulation x virus interaction (F_1,18_ = 0.51, p = 0.49; Figure 4B) and no significant main effect of stimulation (F_1,18_ = 0.33, p = 0.58) or virus (F_1,18_ = 0.065, p = 0.80). All groups successfully discriminated above chance level (eYFP-OFF: t_10_ = 4.30, p = 0.0016; eYFP-ON: t_10_ = 7.62, p = 0.000018; eARCH3.0-OFF: t_8_ = 4.81, p = 0.0010; eARCH3.0-ON: t_8_ = 6.79, p = 0.00014).

Analysis of sample phase object exploration revealed no significant stimulation x virus x sample phase (F_1,18_ = 1.055, p = 0.32); or two-way interactions (stimulation x virus F_1,18_ = 0.65, p = 0.43; stimulation x sample phase F_1,18_ = 0.57, p = 0.46; virus x sample phase F_1,18_ = 0.31, p = 0.59). There was no significant main effect of stimulation (F_1,18_ = 2.38, p = 0.14) or virus (F_1,18_ = 0.13, p = 0.72), but there was a significant main effect of sample phase (F_1,18_ = 5.63, p = 0.029) which reflected greater exploration in sample phase one across all virus and stimulation conditions. Analysis of test phase exploration revealed no significant stimulation x virus (F_1,18_ = 0.12, p = 0.73) and no main effect of stimulation (F_1,18_ = 0.0061, p = 0.94) or virus (F_1,18_ = 0.60, p = 0.45, Table 4). Thus, unlike for OiP and OiC memory, mPFC input to LEC is not required for successful novel object recognition memory.

## Discussion

This study reveals a number of key findings. First, we showed that the mPFC and LEC are independently involved in both the encoding and retrieval of OiP memory following a 1h delay. Second, we demonstrated that encoding and retrieval are dependent on a functional interaction between these regions. Third, activity within the direct mPFC→LEC pathway, during both encoding and retrieval was critical to support longer-term (1 h) but not shorter term (5 min) OiP and an identical pattern of results was revealed when we tested the animals in an OiC task, indicating a consistent role for mPFC→LEC transmission for longer-term associative recognition memory. Finally, as neither pharmacological or optogenetic inhibition impaired novel object recognition we are confident that the observed memory deficits are not a result of generalised cognitive or behavioural impairments. We did find that in some limited examples there were group differences in object exploration. However, it was considered that these are unlikely to have impacted the results. For example, in the disconnection study, simultaneous infusion into the mPFC and LEC prior to the sample phase resulted in a difference in exploration between the CONTRA and IPSI groups which just missed significance. However, from the mean values it is clear that the CONTRA group explore the objects more than the IPSI group so this difference in exploration is unlikely to explain the performance deficit in the CONTRA group.

The finding that the mPFC and LEC, both independently and as a circuit, are critical for different forms of associative recognition memory in rodents is consistent with a number of studies (Barker et al., 2007; Chao et al., 2016; Kuruvilla et al., 2020; Vandrey et al., 2020; Tozzi et al., 2024). We have extended these previous reports providing evidence that the mPFC and LEC are co-involved in both encoding and retrieval. Thus, the OiP memory representations are likely set up within the LEC-mPFC circuit and reactivated for retrieval. While the present experiments clearly do not allow us to delineate the precise roles of the mPFC and LEC within the circuit, other studies have shown that both the mPFC and LEC contain populations of neurons that respond to objects in specific locations (Kim et al., 2011; Hyman et al., 2012; Weible et al., 2012; Morici et al., 2022; Huang et al., 2023), which might suggest that both regions hold specific OiP memory representations. However, the object-place neurons in the mPFC appear to operate over relatively short timescales, while other studies clearly show that the LEC provides the object-place information (Wilson et al., 2013a; Chao et al., 2016) and that object-place neuronal ensembles in the LEC constitute an OiP memory engram (Tozzi et al., 2024). The mPFC may alternatively have a role in the strengthening of associations between object and place and object and context. We have previously shown that retention of OiP across delays of 1 h is dependent on synaptic plasticity processes in the mPFC (Barker and Warburton, 2008). The enhanced synaptic strength in the mPFC may thus be crucial for retaining OiP for 1 h or longer. As such the mPFC and LEC co-dependently operate to support the creation and retention of OiP memory representations, but further work is required to clarify the specific role of the mPFC.

By using optogenetic inactivation methods, which provide directional and temporal control, we were able to examine the importance of the direct mPFC input to the LEC. The original prediction was that inhibition of this projection would impair memory retrieval selectively, based on evidence that the mPFC and outputs to the thalamus, and hippocampus are critical for directing the retrieval of contextual or associative recognition memories (Navawongse and Eichenbaum, 2013; Preston and Eichenbaum, 2013; Barker et al., 2021; Culleton et al., unpublished). However contrary to our prediction we found that the mPFC-LEC projection was also necessary for encoding, all be it following a longer retention delay. The mPFC projection to the LEC terminates in the deeper layers (L5/6) and cells in these deep layers also receive input from the hippocampus, while cells in the superficial layers (L2/3) project to the hippocampus. There is evidence that the LEC and hippocampus work in concert during OiP-like tasks (Chao et al., 2016) although this study did not look at encoding separately from retrieval. Further fan cells in layer 2 of LEC have been suggested to be specifically involved in encoding and forming associations between object, place, and context information, but not simpler object-context association (Vandrey et al., 2020). However, there is also local connectivity between different layers of the cortex (Ohara et al., 2021) and evidence that neurons in the deep and superficial layers show only small differences in spatial processing (Wang et al., 2023). Thus it is possible that OiP and OiC encoding is supported by a mPFC-LEC-CA1 pathway via both the deep and superficial layers of LEC.

The optogenetic experiments also revealed the delay dependence of the necessity of the mPFC→LEC projection, such that mPFC output was necessary for OiP and OiC following the 1 h delay, but not following a 5 min delay. Thus across shorter timescales the mPFC→LEC projection is not critical to support OiP or OiC memory, for which there are at least two possible explanations; either each region can maintain the necessary memory representations within the local circuitry, or that alternative routes of communication between the two areas are more important. Concerning the latter suggestion, there is a direction projection from the LEC to the mPFC (Jones and Witter, 2007; Agster and Burwell, 2009), and there are indirect routes via the hippocampus and nucleus reuniens (Dolleman-van der Weel et al., 2017; Dolleman-van der Weel et al., 2019). It is worth noting that for OiP memory, the NRe appears not to be recruited across shorter timescales (Barker and Warburton, 2018) hence this region is unlikely to be involved. In contrast we have shown that CA1 input to the mPFC is necessary to support shorter term OiP memory (Barker et al., 2021). No studies to date have looked at the role of the LEC→mPFC projection in short-or long-term associative recognition memory, although a recent study showed that inhibition of LEC neurons disrupted mPFC representations of novel items and prevented the formation of new item-outcome associations (Jun et al., 2024).

In summary, pharmacological disconnection of mPFC and LEC shows that mPFC-LEC interaction is required for both memory encoding and retrieval. Optogenetic inhibition of projections from mPFC to LEC showed that this projection was crucial for both encoding and retrieval of both object in place and object in context recognition memory when a 1 h, but not a 5 min, memory retention delay was used. These data show that the direct connection from mPFC to LEC is critical for associative recognition memory, in a delay-dependent manner.

## Acknowledgements

This work was supported by a Wellcome Trust Joint Investigator Award to E.C.W. and Z.I.B. (206401/Z/17/Z), a BBSRC project grant to Z.I.B., E.C.W and P.J.B. (BB/X000915/1) and a BBSRC project grant to E.C.W and G.R.I.B. (BB/Y006402/1).

## Declaration of Interests

The authors declare no competing interests. Generative AI was not used for any aspect of this manuscript.

## Notes

### Competing Interest Statement

The authors have declared no competing interest.

